# *Mycobacterium tuberculosis* exploits SIRT2 for iron acquisition to facilitate its intracellular survival

**DOI:** 10.1101/2024.01.05.574348

**Authors:** Sharmila Talukdar, Radheshyam Modanwal, Gaurav Kumar Chaubey, Asmita Dhiman, Rahul Dilawari, Chaaya Iyengar Raje, Manoj Raje

## Abstract

Iron availability is a critical factor for both bacteria and humans, and its availability significantly influences host-pathogen dynamics. As *Mtb* has coevolved with the human race, *Mtb* relentlessly tries to exploit iron from the tightly regulated iron machinery of host. Sirtuins are evolutionary conserved NAD^+^-dependent deacetylases involved in various cellular processes including infection. Notably, the cytosolic protein, Sirtuin 2 regulates cellular iron homeostasis in hepatocytes and after *Mtb* infection, SIRT2 translocates to the nucleus leading to decreased protective immune response. However, the underlying mechanism as to how *Mtb* exploits SIRT2 for iron acquisition remains unknown. In the current study, we observe that the decreased bacillary load in SIRT2 inhibited or knock down cells is due to low availability of iron to the bacilli. Inhibition or knockdown of SIRT2 in *Mtb* infected cells displays differential modulation of iron import and export proteins suggesting ongoing tussle by host to limit the bioavailability of iron to pathogen. More specifically, by flow cytometry analysis, we show significant upregulation of cell surface Apo Tf and GAPDH in infected SIRT2 inhibited macrophages. Thus, in SIRT2 depleted state, we delineate a different mechanism of iron export occurring through Apo Tf and GAPDH during infection in contrast to the classical iron exporter Fpn1. Collectively, our findings showed the importance of SIRT2-mediated iron regulation in *Mtb* pathogenesis and can encourage designing of novel host-targeted therapeutics.

## INTRODUCTION

Numerous cellular proteins require iron for its optimal functioning. Iron deficiency results in impairment of normal functioning of proteins leading to anaemia or cell death, whereas excess iron and free iron cause cellular injury. Therefore, iron homeostasis within the cell is quintessential and it is achieved by a coordinated regulation of iron storage, uptake and export proteins. To limit the bioavailability of iron, most of the extracellular iron is sequestered by host iron carrier proteins, transferrin (Tf) and lactoferrin (Lf) [1]. Also, upon infection, iron moves from serum to macrophages in reticuloendothelial system to be sequestered into ferritin (iron storage molecule). Tuberculosis caused by the tubercle bacilli, *Mycobacterium tuberculosis* (*Mtb*) accounts for a high burden of global morbidity and mortality[2] [3]. Despite the stringent host defense, *Mtb* has evolved as a remarkably successful intracellular pathogen to survive and multiply within the phagosomes of macrophages and also to persist in host tissues for decades without causing any disease [4]. TB pathophysiology is interconnected with iron metabolism [5]. *Mtb* utilizes a plethora of mechanisms to subvert the host defenses and hijack cellular micronutrient acquisition pathways for its own survival. These include modulation of host metabolic genes and appropriation of host iron [6]. However, little is known about the precise mechanism by which *M. tuberculosis* continues to ensure adequate iron delivery within the macrophage phagosome.

Earlier our group has shown that the cytosolic NAD^+^-dependent glycolytic enzyme Glyceraldehyde-3-phosphate dehydrogenase (GAPDH) acts as a non-canonical receptor for the mammalian iron carrier proteins Transferrin (Tf) and Lactoferrin (Lf) [7] [8]. GAPDH has also been reported to be secreted into the extracellular millieu for iron delivery [9,10]. While the canonical function of GAPDH had been described as a metabolic enzyme with a key role in glycolysis, however numerous subsequent studies have demonstrated that it performs multiple unrelated functions within the cell and in the extracellular milieu. The class III histone deacetylase (HDAC) members, called sirtuins, have different subcellular localizations to carry out their respective cellular processes [11]. The mechanisms by which different pathogenic bacteria exploit host SIRT2 for its survival within the cell have been explored in recent years [12–14]. Among other sirtuins, SIRT2 is the only sirtuin that resides predominantly in the cytoplasm. This cellular localization of SIRT2 is consistent with previously identified cytosolic iron–sensing systems. An elegant study earlier demonstrated that iron is regulated by Sirtuin 2 with the help of iron export protein, ferroportin1 [15]. Increasing evidence is unfolding the influence of pathogens on host proteins. However, our knowledge on the interplay of host proteins in driving the pathogenesis of the tricky pathogen, *Mycobacterium tuberculosis* is in its infancy.

In this study, we reveal a central role of host SIRT2-mediated iron regulatory response upon infection of *Mtb*. We found that there is a differential modulation of iron import and export proteins upon pharmacological inhibition of SIRT2/ lentiviral or siRNA mediated SIRT2 knock down in infected and uninfected conditions. We observed that SIRT2 inhibition restricts *Mtb* growth in primary and secondary macrophages. One of the reasons for reduced *Mtb* survival in the SIRT2 KD or inhibited macrophages was due to low iron content inside macrophages. Interestingly, upon iron supplementation, growth of *Mtb* was observed to be significantly rescued in SIRT2 KD or AGK2 treated cells. To maintain iron homeostasis, cells need to maintain a fine balance between their influx and egress out of cells. In this study, we observed an increase in apo transferrin receptor and GAPDH in SIRT2 KD cells after *Mtb* infection in comparison to Wildtype (WT) or Empty vector (EV) cells. These alterations could be a manifestation of the cellular homeostatic and immune machinery attempting to limit the availability of intracellular iron so as to restrict the replication of intracellular pathogen.

## RESULTS

### SIRT2 regulates macrophage iron content

SIRT2 has been reported to regulate iron in mouse embryonic fibroblasts (MEFs) and human hepatocyte (HepG2) cells [15]. Fe is an essential cofactor for 40 different enzymes encoded within the *Mtb* genome and hence *Mtb* bacilli demands high iron [21]. There is a stringent regulation of iron by the different proteins of iron regulatory machinery inside the host to restrict iron in response to *Mtb* infection. To address the importance of SIRT2-mediated iron regulation in *Mtb* pathogenesis, we began with assessing whether SIRT2 modulates iron in immune sentinel, macrophages. Three principle approaches were employed to deplete the cellular SIRT2 content: lentiviral mediated SIRT2 knock down, transient transfection with SIRT2 siRNA and the well-studied SIRT2 inhibitor (AGK2) [22]. The knock down of SIRT2 in THP1 has been confirmed by both Western blot (**Fig. 1A)** and RT-PCR **(Fig. S1A**). We first evaluated the labile iron pool (LIP) in Empty vector (EV) and SIRT2 KD THP-1 cells utilizing calcein-quenching assay. A significantly greater quenching of calcein fluorescence signal was observed in SIRT2 KD THP1 cells as well as in AGK2-treated macrophages when compared to Empty vector or vehicle control cells (**Fig. 1B, 1C**). We next observed the status of ferritin (intracellular iron storage protein) in Empty vector/SIRT2 KD THP1 cells and observed decreased expression of ferritin in SIRT2 KD as compared to Empty vector (**Fig. 1D**). Moreover, the total iron and ^55^Fe content was also found to be decreased upon silencing of SIRT2 or treatment of AGK2 in THP1 cells (**Fig. 1E, 1F**) In light of these above findings, we concluded that SIRT2 has a role in regulating iron levels in immune cells and hints at the importance of this regulator in pathogenesis.

**Fig. 1.**
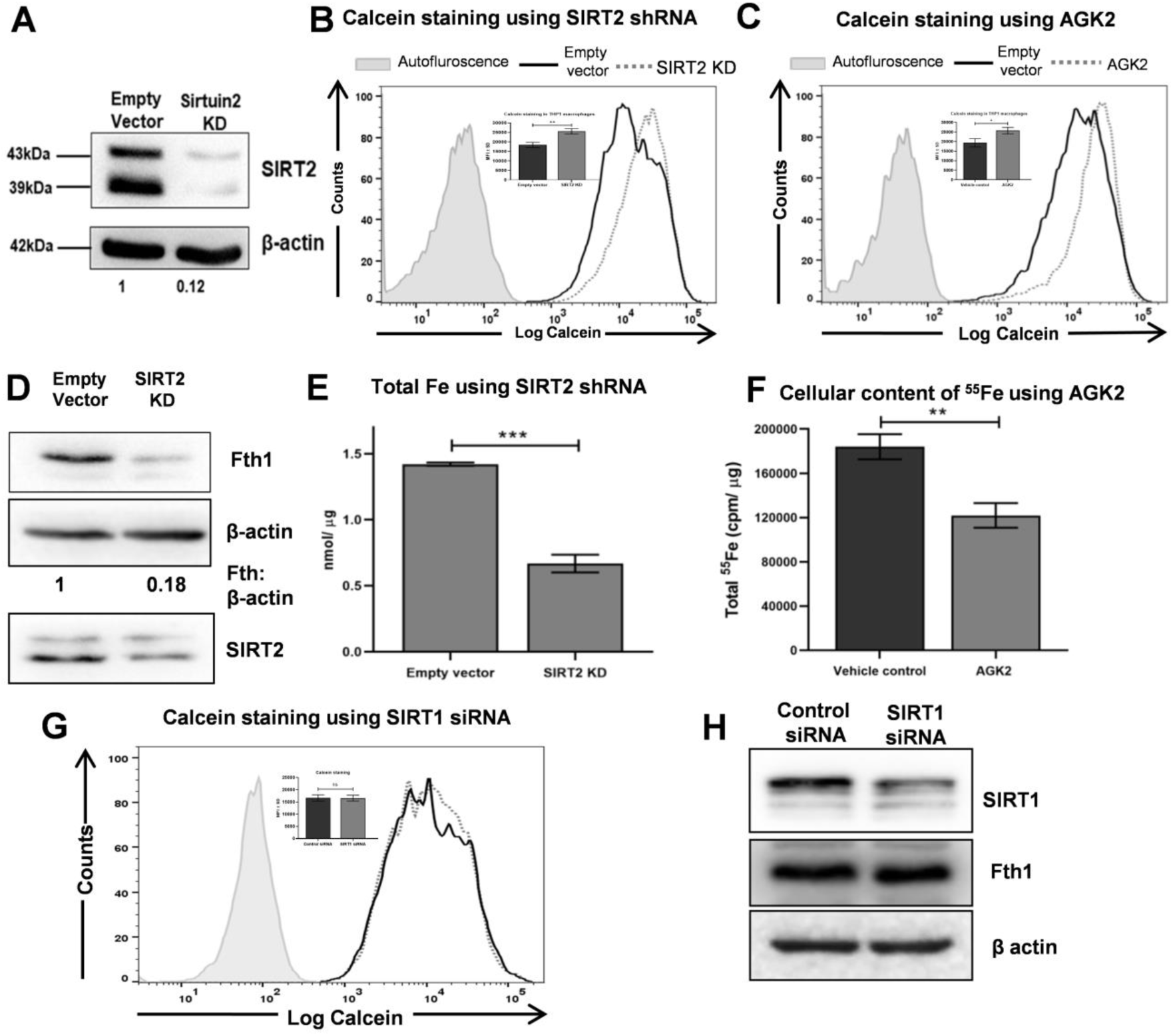
SIRT2 regulates intracellular iron content in macrophages. **(A)** Using Empty vector and SIRT2 lentiviral shRNA particles, SIRT2 KD was confirmed by western blot analysis in THP1 macrophages. Representative blot from 3 independent experiments has been shown **(B)** Assessment of intracellular Calcein-AM staining by flow cytometry in PMA-differentiated Emfpty vector (EV) and SIRT2 KD macrophages **(C)** Intracellular labile iron pool detection in Vehicle control and AGK2-treated THP1 macrophages by Calcein-quenching assay. The calcein fluorescence reflecting labile iron pool was significantly higher in SIRT2 KD/ AGK2 treated macrophages when compared to Empty vector/ Vehicle control cells. Calcein-AM fluorescence is quenched upon binding to intracellular iron and is inversely correlated with intracellular iron content. In SIRT2 KD/ AGK2-treated macrophages, calcein fluorescence is higher due to lesser intracellular iron. Representative flow cytometry overlay has been shown, data in inset are presented as representative plot of background corrected mean fluorescence intensity (MFI ± SD) from 3 independent experiments performed in triplicates (n=3, *p<0.05). **(D)** Representative immunoblot from total cell lysate of Empty vector (EV) and SIRT2 KD THP1 cells for checking the expression of Ferritin (Fth1). Images were quantified by ImageJ software. The expression of Fth1 was found to be significantly decreased as compared to EV cells **(E)** Total iron was estimated from EV and SIRT2 THP1 macrophages and it was found to be lower in SIRT2 KD macrophages in comparison to EV cells (***p < 0.001, n=3) **(F)** PMA-activated THP1 macrophages were incubated for 24h with radioactive iron followed by treatment with DMSO and AGK2 for 24h. The ^55^Fe measured in AGK2-treated cells was decreased in comparison to Vehicle control (DMSO) treated cells (**p < 0.01, n=3) **(G)** Calcein quenching in SIRT1-siRNA transfected macrophages was unchanged as compared to Control siRNA transfected macrophages. Representative flow cytometry overlay has been shown, data in inset are presented as representative plot of background corrected mean fluorescence intensity (MFI ± SD) from 2 independent experiments performed in triplicates (n=3, ns: non-significant) **(H)** Representative immunoblot of SIRT1 siRNA transfected macrophages showed non-significant change in the expression of Fth1 as compared to Control siRNA transfected macrophages.

Among the seven members of Sirtuin family, SIRT1 and SIRT2 share the same compartment inside a cell. Although the predominant localisation of SIRT1 and SIRT2 are nucleus and cytosol respectively, these two proteins shuttle between the nucleus and the cytoplasm under different cellular conditions [23] . Therefore, we were interested to understand whether SIRT1 also regulates intracellular iron. Remarkably, we did not observe iron regulation by SIRT1 as analysed by flow cytometry-based calcein staining and Fth1 expression by Western blot. This shows the specificity of SIRT2-mediated iron regulation in macrophages **(Fig. 1G, 1H)**.

### SIRT2 modulates the expression of surface iron regulatory proteins

Our studies have suggested that SIRT2 is essential for intracellular iron accumulation in macrophages. We next explored the status of the surface receptors involved in iron export and import essential for maintaining iron homeostasis. It has been demonstrated that gene expression of the cellular iron import protein, transferrin receptor (CD71) was significantly higher in Sirt2^−/−^ MEFs than in Sirt2 ^+/+^ MEFs. Also, the cellular iron export protein Fpn1 was noted to be significantly higher at both the mRNA and protein levels in Sirt2 ^−/−^ MEFs [15]. We investigated if similar regulation is evident in macrophages and interestingly, we exhibit an increased surface expression of Fpn1 **(Fig. 2A, 2B)** and CD71 upon SIRT2 KD or inhibition **(Fig. 3A, 3B)**. This result is in accordance with an increased binding of the Holo Tf in both AGK2-treated and SIRT2 KD macrophages **(Fig. 3C, 3D).** The increased expression of surface CD71 in SIRT2 KD or inhibited cells can be explained by the fact that the number of TfR1 on the cell surface is regulated based on the amount of intracellular iron according to cell’s need. Therefore, when the intracellular iron levels drop, the cell recruits more TfR1 on its membrane [24]. Further, we were interested to understand whether cells with decreased SIRT2 activity can respond with appropriate modulation of their receptors for iron acquisition in response to perturbations in iron levels. To study this, we subjected Empty vector THP-1 and SIRT2 KD THP-1 cells to iron loading by supplementing the culture medium with 100 µM FeCl_3_. We found that the increase in surface CD71 in SIRT2 KD cells significantly decreases thus demonstrating that external supplementation of iron can still regulate the SIRT2-dependent surface CD71. This suggests that the elevated CD71 observed on the KD cells may be a cellular response occurring as a consequence of cells being depleted of iron (**Fig. 3E and 3F**).

**Fig. 2.**
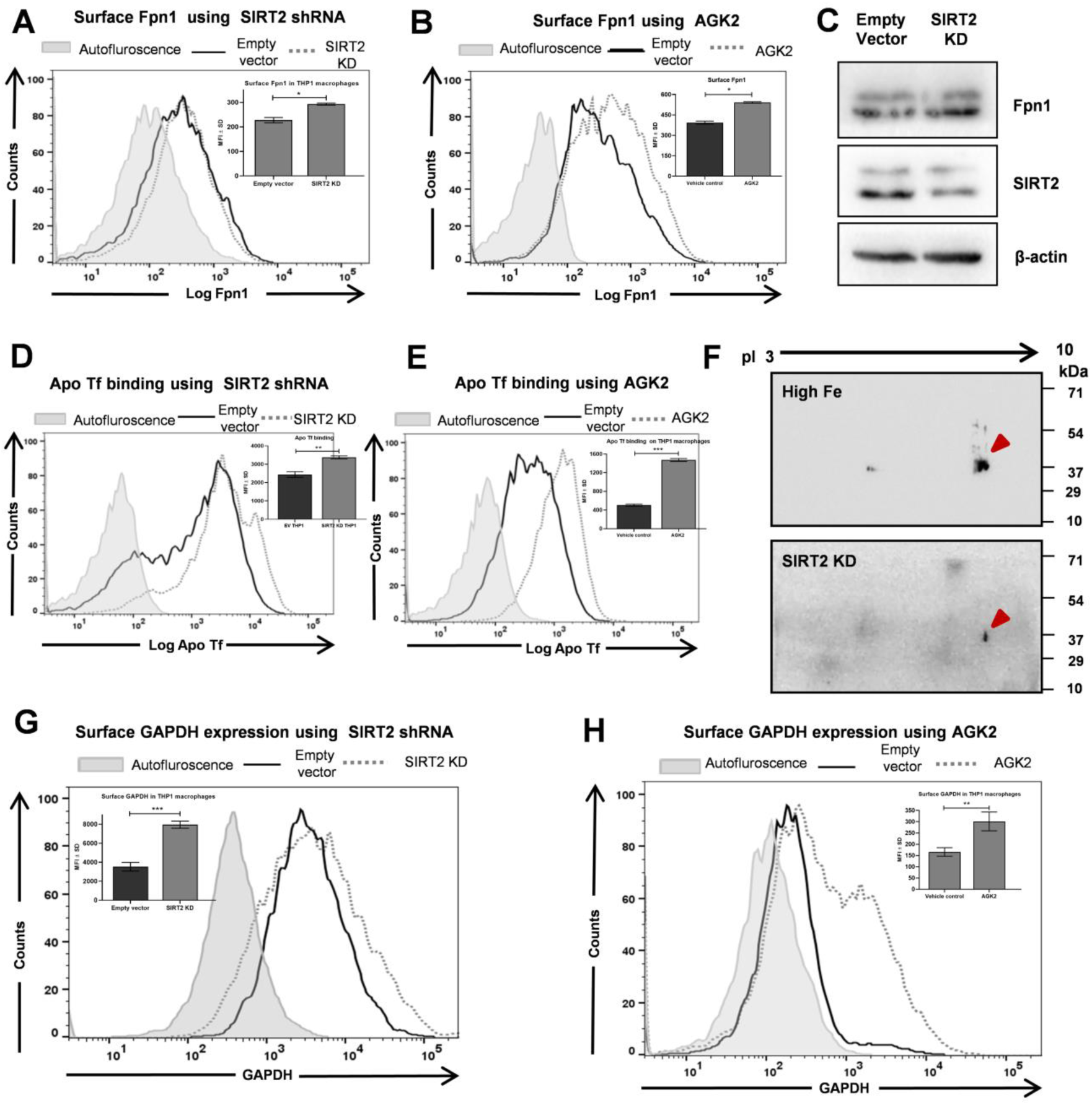
SIRT2 up regulates the expression and binding of classical and the non-classical iron export machinery in macrophages. **(A)** Cell surface expression of Fpn1 in EV and SIRT2 KD macrophages by flow cytometry **(B)** PMA-activated THP1 macrophages were treated with DMSO and 10 µM AGK2 for 24h followed by cell surface expression of Fpn1 by flow cytometry. Representative flow cytometry overlay has been shown, data in inset are presented as representative plot of background corrected mean fluorescence intensity (MFI ± SD) from 3 independent experiments performed in triplicates (n=3, *p<0.05). **(C)** Representative immunoblot of Fpn1 from whole cell lysate of EV and SIRT2 KD THP1 macrophages **(D)** EV THP1 and SIRT2 KD macrophages were incubated with labelled Apo Tf for 1h at 4°C followed by flow cytometry analysis **(E)** Control and AGK2-treated THP1 macrophages incubated with labelled Apo Tf for 1h at 4°C were analysed after data acquisition in flow cytometer. Representative flow cytometry overlay has been shown, data in inset are presented as representative plot of background corrected mean fluorescence intensity (MFI ± SD) from 3 independent experiments performed in triplicates (n=3, **p<0.01, ***p<0.001) (**F)** Western blot representation of 2D-gel-electrophoresis separated GAPDH isoforms from membrane fractions of SIRT2 KD and high iron (100µM)-treated THP1 cells, pI: isoelectric point. Arrows in red depict the alkaline shift or isoelectric point (∼7) in both SIRT2 KD and high iron-treated THP1 cells **(G)** Cell surface GAPDH on EV and SIRT2 KD macrophages was quantified by flow cytometry using anti-GAPDH antibody **(H)** Control and AGK2-treated macrophages were incubated with anti-GAPDH antibody for quantification by flow cytometry . Representative flow cytometry overlay has been shown, data in inset are presented as representative plot of background corrected mean fluorescence intensity (MFI ± SD) from 3 independent experiments performed in triplicates (n=3,***p<0.001).

**Fig. 3.**
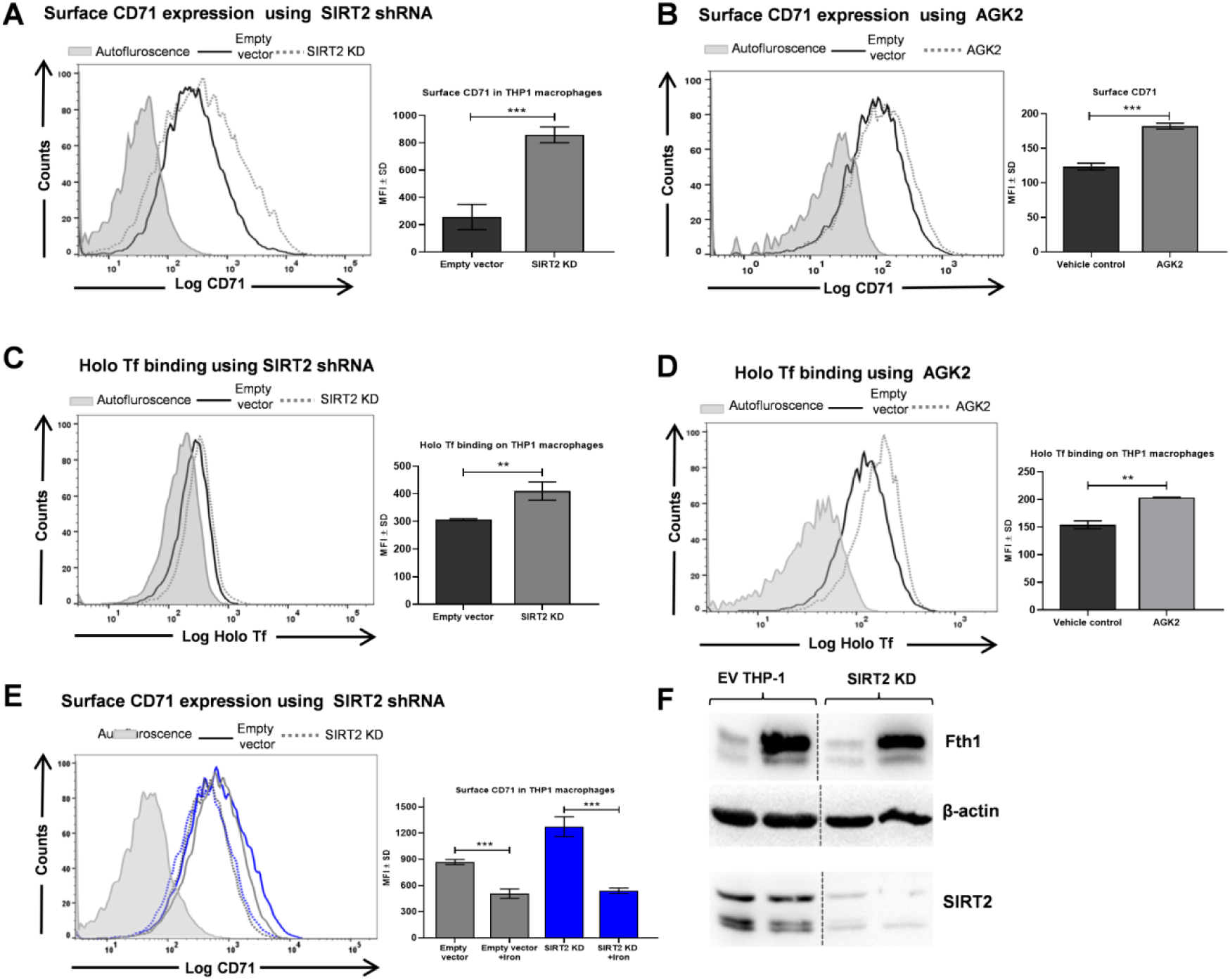
SIRT2 up regulates the classical iron importer, transferrin receptor (CD71) **(A)** PMA-activated and differentiated EV and SIRT2 KD THP1 macrophages were incubated with primary CD71-APC labelled antibody at 4°C for 1h and Cell surface CD71 was quantified using flow cytometry **(B)** THP1 cells after activation and differentiation were treated with 10 µM AGK2 for 24 h and incubated with labelled CD71 antibody for determination by flow cytometry. Data are shown as representative overlay histogram and data in inset are presented as representative plot of background corrected mean fluorescence intensity (MFI ± SD) from 3 independent experiments performed in triplicates (n=3,***p<0.001) **(C-D)** Quantification of labelled Holo Tf binding in EV/ Control and SIRT2 KD/ AGK2-treated macrophages by flow cytometry. Representative flow cytometry overlay has been shown, data in inset are presented as representative plot of background corrected mean fluorescence intensity (MFI ± SD) from 2 independent experiments performed in triplicates (n=3, ***p<0.001) **(E)** EV and SIRT2 KD THP1 macrophages were given external supplementation of high iron (100 µM) for 24h followed by flow cytometric analysis of CD71 using anti-CD71 APC antibody. Data are shown as representative overlay histogram and data in inset are presented as representative plot of background corrected mean fluorescence intensity (MFI ± SD) from 2 independent experiments performed in triplicates (n=3, ***p<0.001) **(F)** Representative western blot to check Fth1 expression from the lysates of high iron treated and untreated EV and SIRT2 KD cells.

Previous study from our group has reported that surface GAPDH is recruited in response to cellular iron loading and functions as a receptor for Apotransferrin (ApoTf) [19]. Also, we did not observe any significant difference in the Fpn1 expression from total cell lysate of EV and SIRT2 KD cells **(Fig. 2C)**. Therefore, we were curious to decipher the role of GAPDH and Apotransferrin (ApoTf) in SIRT2 KD model system and interestingly we observed a significant up-regulation of both surface Apo Tf binding **(Fig. 2D, 2E)** and GAPDH upon the depletion of SIRT2 or SIRT2 activity (**Fig. 2G, 2H** ). This observation was also investigated in a murine macrophage model. To this end, SIRT2 KD J774 macrophages were prepared using SIRT2 specific lentiviral particles and SIRT2 KD was confirmed by Western blot **(Fig. S2A)**. Similar to the result obtained in THP1 macrophages, an increase in Apo Tf binding and surface GAPDH expression was consistent in SIRT2 KD and AGK2-treated J774 macrophages in comparison to EV and control macrophages **(Fig. S2B-E)**.

Further, we performed an *in vivo* pull-down assay to examine protein-protein interaction of GAPDH and Apo Tf from the cell membranes. Affinity pull down of GAPDH with biotinylated apo-transferrin from the membrane fractions indicated the interaction is prevalent in SIRT2 depleted state **(Fig. S2G)**. Further, investigations utilizing confocal microscopy revealed a distinct colocalization between GAPDH and Apo Tf on the plasma membrane. Quantification by Pearson’s correlation coefficient showed an increased colocalization of GAPDH and Apo Tf in SIRT2 KD macrophages compared to EV cells **(Fig. 6A)**.

### GAPDH recruited on the membrane surface is an alkaline isoform

There are numerous reports on the existence of different isoforms of GAPDH present across all species available in literature. This is due to a plethora of posttranslational modifications that this protein is known to undergo. Specific isoforms of GAPDH have been known to be selectively recruited to different cellular compartments when cells undergo certain types of stress [25]. In yeast, after H_2_O_2_ treatment, a new form of GAPDH (Tdh3p) with a lower isoelectric point has been reported in mitochondrial extracts showing the occurrence of a posttranslational modification [26]. An elegant study earlier reported that when cells are subjected to the stress of iron overloading there occurs a switch in the isoform of GAPDH recruited onto the cell membrane wherein it functions as a receptor for apo-transferrin (iron free form of transferrin) and functions to remove excess iron from cells [19]. In a different study, it has been observed that unlike the GAPDH recruited in high iron condition, the GAPDH exposed during apoptosis revealed an acidic shift in 2D-PAGE analysis [27]. Thus, we assessed the nature of GAPDH isoform recruited to the plasma membrane of macrophages deficient in SIRT2. For this the membrane fraction of SIRT2 KD THP-1 cells were purified and subjected to two-dimensional (2D) gel electrophoresis followed by immunoblot for detection of GAPDH and compared with high iron (100µM) –treated THP1 macrophages. **(Fig. 2F)**. Strikingly, we observed that the GAPDH revealed an alkaline shift with a pI of ∼7 in the plasma membrane fraction of SIRT2 KD macrophages. Moreover, high iron treated cells also showed the same shift of pI ∼7. Probably, the same isoform of GAPDH is playing a role in exporting iron under SIRT2 depletion as well as high iron condition.

### SIRT2 inhibition restricts *Mtb* growth in an iron-dependent manner

To delineate the physiological role of SIRT2 in intracellular survival of *Mtb*, survival of the bacteria was assessed in Empty vector (EV) and SIRT2 KD THP1 macrophages by plating the cell lysate at 0h, 24h and 48h post-infection. In SIRT2 KD cells, the bacteria showed decreased survival as compared to EV cells. This is further validated by a decreased survival of intracellular bacteria in AGK2-treated bone-marrow derived-macrophages in comparison to untreated macrophages **(Fig. 4A, 4B)**. It has been reported that in AK7 (a SIRT2 inhibitor) treated dendritic cells, *Salmonella Typhimurium* showed an increased survival as compared to control [13]. However, in our case, enumeration of intracellular *Mtb* suggested that the capability of SIRT2 to increase intracellular mycobacterial growth is attenuated upon SIRT2 KD or SIRT2 activity inhibition. We hypothesized that the decreased bacillary load might be due to the lower availability of iron in AGK2 treated/ SIRT2 KD cells. Therefore, upon iron supplementation in AGK2 treated/ SIRT2 KD cells, an enhanced survival of bacteria was observed thus rescuing the intracellular bacilli from the iron deprived environment created by SIRT2 depletion. It has been well known that Fe supplementation helps in *Mtb* proliferation and Fe starvation hampers *Mtb*’s ability to proliferate [3]. As SIRT2 KD or AGK2-treated cells are already deprived of iron, the observed increase in bacillary load after Fe supplementation in SIRT2 KD or AGK2-treated cells was less than Fe-supplemented Empty vector or untreated cells. These findings indicate that SIRT2-mediated iron regulation and control of mycobacterial growth depends on SIRT2 **(Fig. 4A, 4B).** Similar pattern of bacillary load was observed in infected macrophages incubated for 48h **(Fig. S3A and S3B)**. It is important to note that *Mtb* infection leads to an increased expression of SIRT2 at both mRNA and protein level **(Fig. S3D and S3E)** and this observation is in accordance with a previous report [12]. In agreement with our earlier observations, SIRT2 positively regulates intracellular iron, and hence up regulation of SIRT2 is beneficial for the *Mtb* residing inside the host to get a copious amount of iron.

**Fig. 4.**
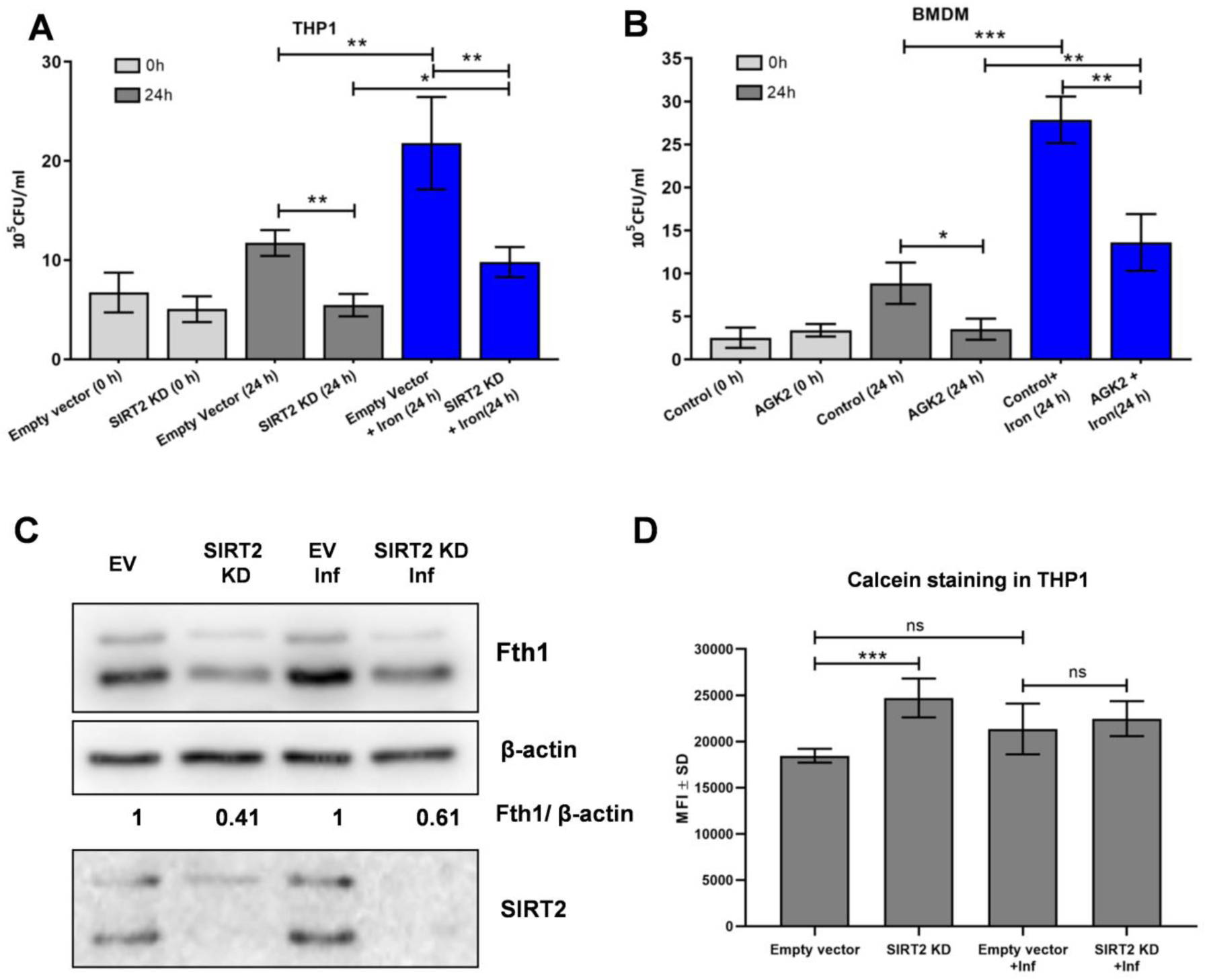
SIRT2 aids in mycobacterial survival inside macrophages in an iron-dependent manner. **(A)** Empty vector and SIRT2 KD THP-1 macrophages were infected with 1:10 MOI of *Mtb* H37Rv and 100 µM FeCl_3_ was added wherever appropriate. Cells were processed for CFU enumeration at 0 h and 24 h hours post infection. Results are indicated as 10^5^ CFU/mL (***p<0.0001 and **p<0.01, *p<0.05, n=3). Data represents average of two independent experiments performed in triplicates **(B)** Bone marrow derived macrophages were treated with or without AGK2 for 24 h followed by infection with 1:10 MOI of *Mtb* H37Rv and 100 µM FeCl_3_ wherever appropriate. After certain time points, CFU load was measured. Results are indicated as 10^5^ CFU/mL (***p<0.0001 and **p<0.01, *p<0.05, n=3). Data represents average of two independent experiments performed in triplicates **(C)** Representative blot showing expression of Fth1 with or without H37Rv infection in EV and SIRT2 KD THP1 macrophages **(D)** Measurement of intracellular calcein staining reflecting labile iron pool in uninfected and infected EV and SIRT2 KD THP1 cells. Data is representative of 2 independent experiments performed in triplicates (n=3, ***p<0.001, ns non-significant).

### *Mtb* infection fails to increase intracellular Fe in SIRT2 depleted macrophages

Within the host, *Mtb* infection leads to the sequestration of iron in iron-storage proteins such as ferritin in order to limit Fe access to the intracellular bacilli. SIRT2 is an essential player in increasing or maintaining the reservoir of labile, stored and total iron content of macrophages as shown in this study. After infection in Empty vector/ Control siRNA transfected macrophages, the expression of ferritin was significantly increased. Next, we were interested to know the status of ferritin in infected SIRT2 KD/ SIRT2 siRNA transfected macrophages, and interestingly, we observed a decrease in expression of Fth1 similar to uninfected SIRT2 KD/SIRT2 siRNA transfected macrophages **(Fig. 4C, S3C)**. This suggests that in SIRT2-deficient condition, *Mtb* is not able to increase the intracellular iron content (as shown by decreased Fth1 expression), when compared to EV/ Control siRNA transfected cells, as SIRT2-mediated iron regulation is dominant and not *Mtb* infection. Furthermore, labile iron pool of macrophages was evaluated in uninfected/ infected EV and SIRT2 KD macrophages. As expected, though there was an upregulation of labile iron pool in SIRT2 KD macrophages but we observed a non-significant difference in the labile iron pool of infected EV and SIRT2 KD macrophages thus supporting the previous result **(Fig. 4D)**.

### *Mtb* infection results in differential modulation of cell surface iron regulatory proteins

To understand the interplay of the iron exporters and importers in infected scenario, we next examined the expression of major iron responsive proteins: CD71, Fpn1, Apo Tf and GAPDH using flow cytometry analysis wherein we gated the infected and uninfected population. Under infected condition, the expression of the classical iron import receptor, CD71 is generally down regulated as a host defense to limit the import of iron for the invading pathogen [28]. In our case, in infected EV as well as SIRT2 KD THP1 macrophages, a significant decrease in the expression of CD71 was observed compared to uninfected cells as a normal host response to *Mtb* infection **(Fig. 5A)**. Notably, the decrease in CD71 along with a concomitant decrease in binding of its ligand holo-transferrin **(Fig. 5B)** in infected cells was SIRT2 independent. On the other hand, the expression of Fpn1, Apo Tf and GAPDH was increased in infected EV macrophages compared to uninfected EV cells. Importantly, in infected SIRT2 KD macrophages; there was an even more increased expression of the exporters compared to infected EV cells **(Fig. 5C-E)**. Depletion of SIRT2 is leading to egress of iron during infection thus hampering *Mtb* to establish pathogenesis and it suggests an intriguing host defence strategy to limit iron availability to intracellular pathogen. In parallel, total iron in SIRT2 KD and infected SIRT2 KD cells were estimated to be significantly lower because of the active expelling of iron via the exporters **(Fig. 5F)**.

**Fig. 5.**
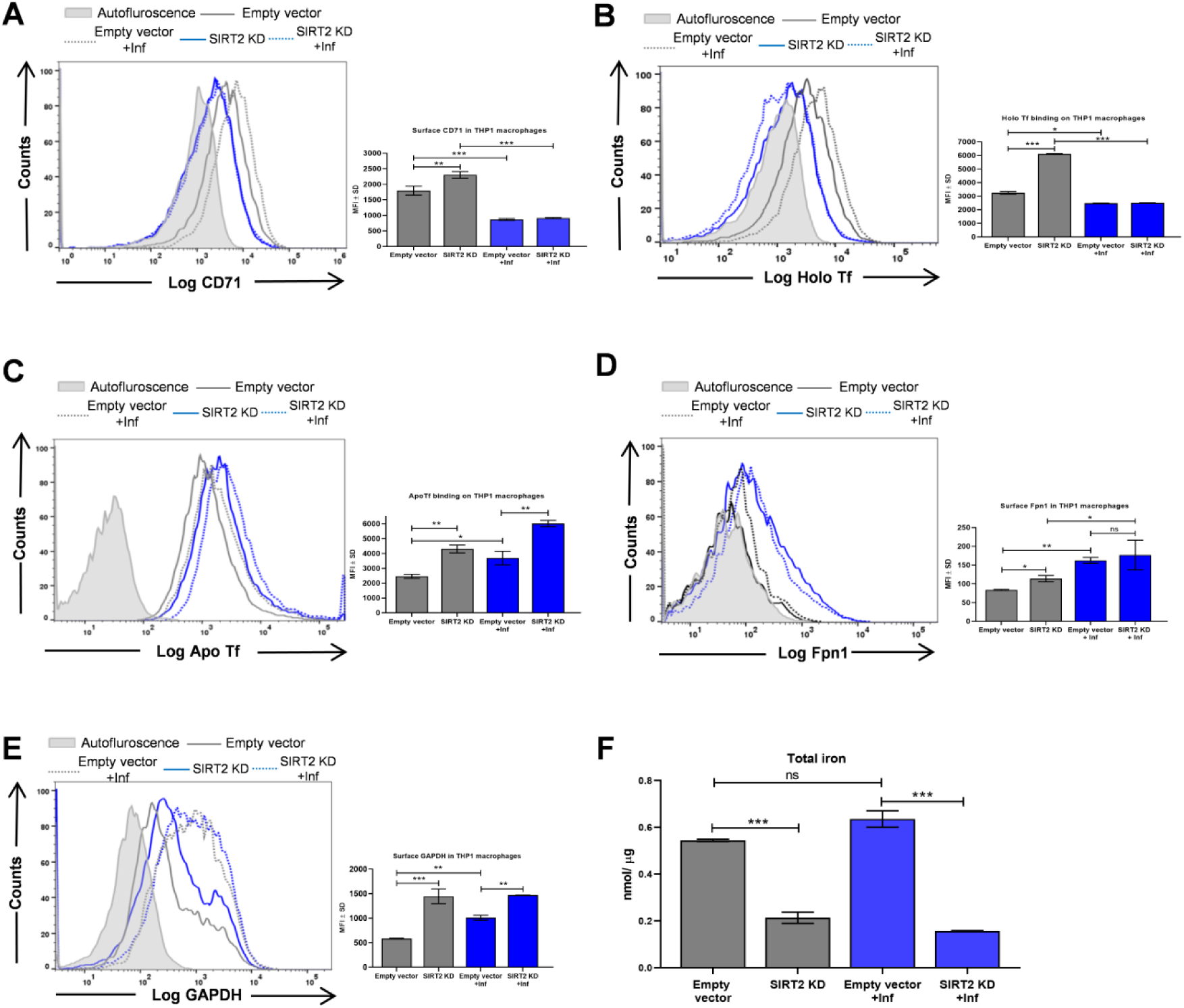
Differential modulation of iron responsive machinery by SIRT2 upon *Mtb* infection. **(A)** PMA-activated and differentiated EV and SIRT2 KD THP1 macrophages were infected with or without *Mtb* at 1: 10 MOI for 24 h. The cells were then incubated with primary CD71-APC labelled antibody at 4°C for 1h and cell surface CD71 expression was quantified using flow cytometry **(B)** Quantification of labelled Holo Tf binding in uninfected or infected EV and SIRT2 KD macrophages by flow cytometry. Representative flow cytometry overlay has been shown for the above figures, data in inset are presented as representative plot of background corrected mean fluorescence intensity (MFI ± SD) from 2 independent experiments performed in triplicates (n=3, ***p<0.001) **(C)** Binding of Apo Tf in uninfected or infected EV and SIRT2 KD macrophages was assessed after incubation of Apo Tf-Alexa Fluor 647 at 4°C for 1h **(D)** Cell surface expression of Fpn1 in uninfected or infected EV and SIRT2 KD macrophages by flow cytometry **(E)** Uninfected or infected EV and SIRT2 KD macrophages were incubated with anti-GAPDH antibody for quantification of cell surface GAPDH by flow cytometry. Representative flow cytometry overlay has been shown, data in inset are presented as representative plot of background corrected mean fluorescence intensity (MFI ± SD) from 2 independent experiments performed in triplicates (n=3, ***p<0.001) **(F)** Measurement of total iron content in EV and SIRT2 KD macrophages under uninfected and infected conditions. Quantified data has been depicted in nmol/ µg. Statistical significance was done using unpaired t-test (n=2, ***p<0.001). Data represents one of the two independent experiments performed in duplicates.

These findings suggest that in response to *Mtb* infection in SIRT2-deficient cells, the host down regulates the classical iron import receptor, CD71 to limit the iron availability to the invading pathogen. However, it upregulates the molecules involved in iron export viz. Apo Tf and GAPDH to facilitate export of iron from the host cell.

### SIRT2 regulates GAPDH-mediated egress of iron

To gain more mechanistic insight as to whether GAPDH functions as a receptor for Apo Tf to extrude intracellular iron, the Apo Tf binding and intracellular iron was analysed in GAPDH and SIRT2 siRNA transfected macrophages and compared with only SIRT2 siRNA transfected macrophages. Cells treated with control siRNA were set up in parallel. Interestingly, we observed that upon knockdown of GAPDH and SIRT2, there was less Apo Tf binding and more intracellular accumulation of iron as compared to only SIRT2 knockdown **(Fig. 6C, 6D)**. This suggests that in a SIRT2 depleted cellular environment, GAPDH is vital in increasing Apo Tf binding so as to export the trapped iron. A concentration of 40nM GAPDH and SIRT2 siRNA respectively has been utilized for silencing GAPDH and SIRT2 **(Fig. 6B)**. Surface knock down of GAPDH was confirmed using GAPDH antibody and analyzed by flow cytometry **(Fig S6)**.

**Fig. 6.**
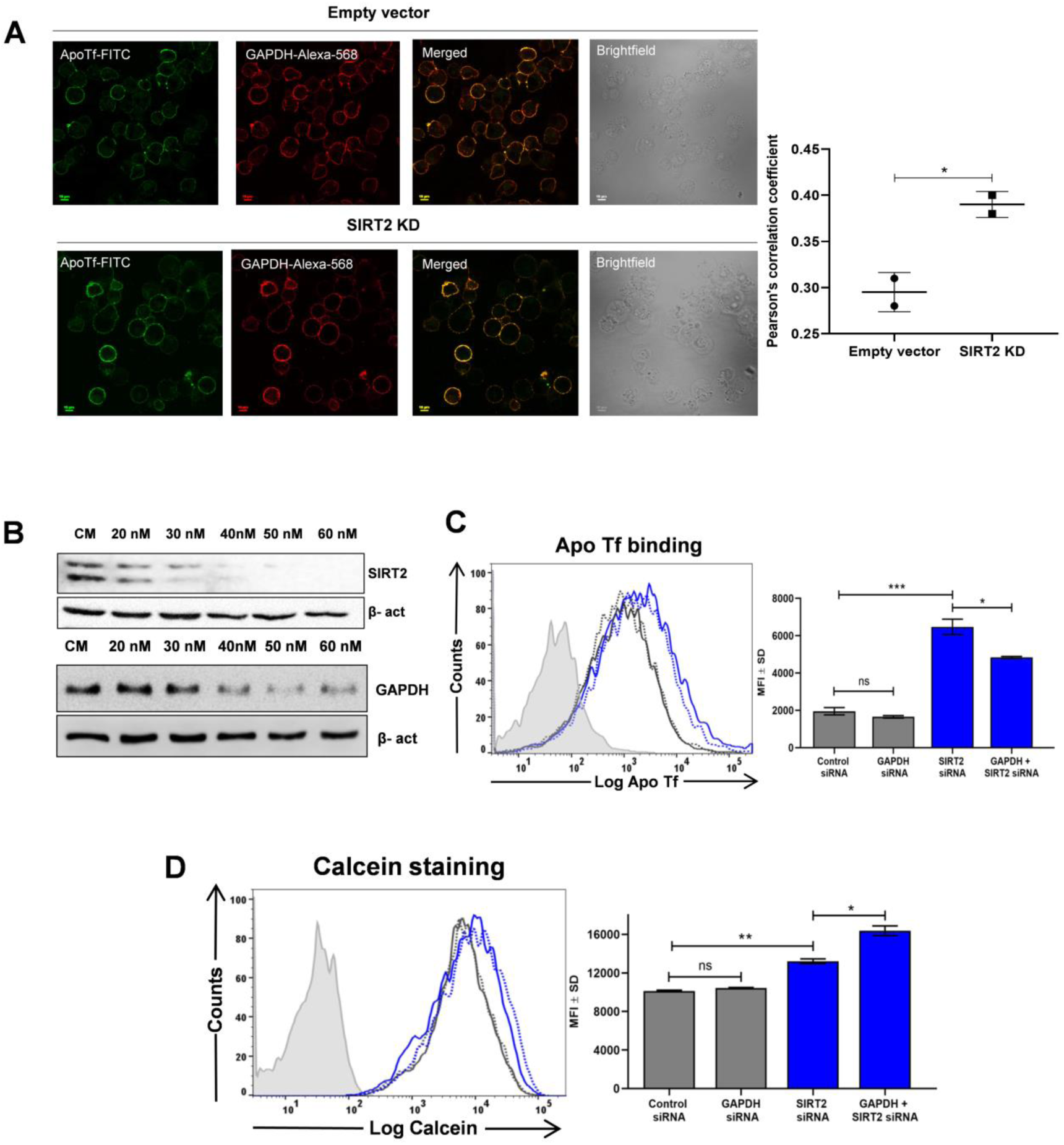
SIRT2 regulates GAPDH-mediated egress of iron. **(A)** Colocalisation of GAPDH and Apo Tf-FITC on the surface of cell membrane in EV and SIRT2 KD THP1macrophages, as visualised using confocal microscopy. Representative image scale bars, 10 µm. Pearson’s correlation coefficient was calculated to evaluate the extent of colocalisation (*p<0.05, n=15) **(B)** Western blot of SIRT2 and GAPDH in THP1 macrophages transfected with SIRT2 or GAPDH-specific siRNA in a dose-dependent manner **(C-D)** THP-1 macrophages were transfected with GAPDH or SIRT2 or GAPDH and SIRT2 siRNA using Lipofectamine 2000 for 48 hours. Cells treated with control siRNA were set up in parallel. Apo Tf binding was evaluated in all the conditions and intracellular calcein staining was checked using flow cytometry. Representative flow cytometry overlay has been shown, data are presented as representative plot of background corrected mean fluorescence intensity (MFI ± SD) from 2 independent experiments performed in triplicates (n=3, ***p<0.001, **p<0.01, *p<0.05, ns non-signficant).

## DISCUSSION

*Mycobacterium tuberculosis* is a highly contagious intracellular human pathogen continuing to evolve numerous drug resistant strains at an alarming rate [29]. At the same time, unscientific and excessive utilization of antibiotics in recent years has increased the emergence of multi drug resistant bacteria [30].

*Mtb* requires copious quantities of iron for its growth and survival in mammalian host and consequently the cellular machinery involved in this process of iron transport to the invading pathogen is an attractive emerging drug target. Recently, the development of anti-TB therapeutic options, that could target these iron acquisition pathways, has been constrained by the dearth of information on *Mtb* iron acquisition [31]. Our understanding of host-pathogen interactions suggests that adding host-directed techniques to the current anti-TB therapy may; boost bacterial clearance, shorten treatment periods, decrease the development of drug-resistant strains, and reduce the likelihood of disease relapse and recurrence.

Host-directed therapy (HDT) for TB treatment is a feasible concept under which host response is targeted, for example, the directed modulation of host inflammatory pathways. Similarly, it is plausible that the iron acquisition mechanisms or molecules exploited by *Mtb* inside the host can also be targeted. Based upon our previous fndings, *Mtb* hijacks mammalian secreted Glyceraldehyde-3-phosphate dehydrogenase (sGAPDH) for the targeted delivery of host iron carrier proteins Lactoferrin (Lf) and Transferrin (Tf) to intracellular bacilli [32].

Sirtuins are a group of evolutionary conserved NAD^+^-dependent enzymes having different biological roles. One such important role of intracellular iron regulation by SIRT2 has been reported in MEF’s and HepG2 cells. It has been found that SIRT2 removes the acetyl group from nuclear factor erythroid-derived 2–related factor 2 (NRF2), which leads to reduced expression of Ferroportin1 (Fpn1) and increase in cellular iron [15]. For its success as an intracellular pathogen, *Mtb* skews multiple host pathways in its favour. Additionally, *Mtb* infection modifies the host’s transcriptional landscape dramatically by secreting a variety of virulence proteins [33]. Till date, very few studies have clearly demonstrated the role of sirtuins in bacterial pathogenesis.

In light of these observations, we were keen to decipher the role of SIRT2-mediated iron exploitation in TB pathogenesis. We proceeded to investigate these using infected macrophage model systems while chemically inhibiting SIRT2. Additionally a SIRT2 knock down cell line was also utilized to decipher its role in iron homeostasis of *Mtb* infected cells. Upon monitoring the iron status of cells we observed a complete decrease in cellular iron. This was due to decrease of both, the intracellular labile as well as stored iron pools of SIRT2 KD or AGK2-treated cells. We also found an increase in the surface expression of CD71 and GAPDH on the surface of these cells. This correlated with an increase of holo Tf binding on SIRT2 KD macrophages. Knockdown of SIRT2 also resulted in a significant upregulation of surface GAPDH. Upon investigation we discovered that the isoform of GAPDH in SIRT2 KD cells had an isoelectric point with an alkaline shift. Previously from our lab, it has been reported that when cells have an excess of intracellular iron, there is a switch of GAPDH on the cell surface to a more basic isoform which binds to Apo-Tf [19]. Matching this observation we too found a significant increase in the binding of apo-Tf to the cells wherein SIRT2 had been depleted.

The importance of SIRT2 in TB pathogenesis has been reported recently wherein, they observed that *Mtb* infection increases the expression of SIRT2 and the movement of SIRT2 to nucleus impacts histone acetylation as well as other immune modulators. Pharmacological inhibition of SIRT2 activity lowers CFU burden in both *in vitro* and *in vivo* model systems. Notably, the widely used SIRT2 inhibitor along with first-line anti-TB drugs was shown to be potent and efficient in defense against highly infectious drug-resistant strains of *Mtb* [12]. Our investigations also revealed an increase in SIRT2 expression after *Mtb* H37Rv infection of peritoneal and THP1 derived macrophages.

Since SIRT2 regulates iron and *Mtb* upregulates SIRT2, we were interested to see if SIRT2 has any effect on the intracellular survival of *Mtb*. We measured the intracellular survival of *Mtb* in BMDMs upon addition of AGK2 (a SIRT2 inhibitor) and also in SIRT2 KD THP1 cells. A significantly decreased bacillary load was observed in cells wherein the activity of SIRT2 had been circumscribed. It is well known that Fe supplementation helps in *Mtb* proliferation and Fe starvation hampers *Mtb*’s ability to proliferate [3]. We hypothesized that the decreased bacillary load could be the outcome of a depleted iron level in AGK2 treated or SIRT2 KD cells. To confirm this hypothesis that lower load of *Mtb* is due to less iron present in the AGK2-treated or SIRT2 compromised cells, we provided iron supplementation to infected SIRT2 KD cells. The result was a significant increase in the survival of intracellular mycobacteria thereby indicating that iron supplementation had rescued the survival of *Mtb*. In comparison to wild type or untreated control cells, the SIRT2 KD/AGK2 cells are iron deprived and accordingly, the increase in bacillary load after Fe treatment in SIRT2 KD or AGK2-treated cells was lower than when wild type untreated cells were supplemented with iron.

We also, found that in response to *Mtb* infection in SIRT2-deficient conditions, host cell down regulates the classical iron import receptor, CD71 to limit the iron availability to the invading pathogen. However down regulation of the proteins involved in iron export, (Fpn1, GAPDH and Apo-Tf receptor) does not take place. Consequently while on one hand import of iron is decreased, the export continues. This results in the cells being left with decreased cellular iron content, a condition detrimental to survival of intracellular pathogen.

Previously, our group has shown the interaction of Apo transferrin with GAPDH on the surface of plasma membrane under excess iron. We were keen to confirm the role of SIRT2 in mediating the export of iron via GAPDH and Apo Tf. Utilizing siRNA mediated transfection; we were able to find a positive correlation of SIRT2 in exploiting GAPDH leading to extrusion of intracellular Fe. A schematic representation of the events has been depicted in **Fig. 7**. From this study, it can be implied that inhibitors of SIRT2 can be used as an adjunct therapy to combat TB pathogenesis and by understanding the mechanistic aspect of host after *Mtb* infection; new avenues in combating TB pathogenesis can be deciphered.

**Fig. 7.**
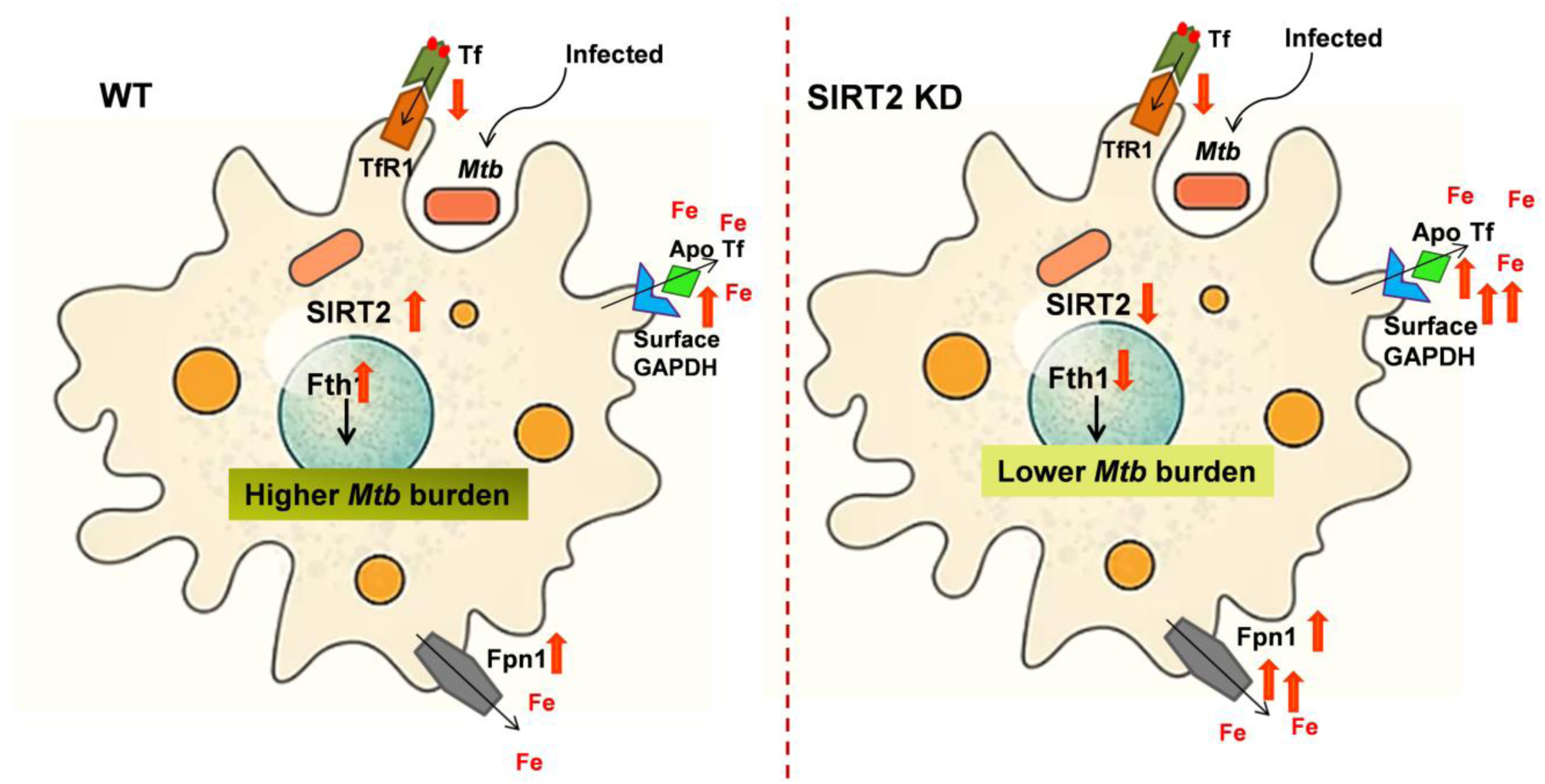
Schematic representation of the tussle between host to restrict the bioavailability of iron and *Mtb*’s attempt of up regulating SIRT2 to acquire iron.

### Materials and methods

#### Mice

C57BL/6 (6–8 weeks of age) male mice were procured from animal facility (iCARE) of CSIR-IMTECH. The animals were maintained in the animal facility of CSIR IMTECH, Chandigarh, India.

#### Bacterial culture

*Mtb* H37Rv and *Mtb* H37Rv expressing GFP were cultured in 7H9 medium (Middlebrook, Difco) with 0.2% glycerol, 0.1% Tween 80 and 10% OADC. GFP-*Mtb* H37Rv was maintained under selection pressure of hygromycin (50 µg/ml) as described earlier [16,17]. Stocks of cultures grown to mid-log phase were preserved in freezing medium (complete 7H9 medium and 20% glycerol) and stored at −80°C. All experiments were carried out using these cryopreserved stocks. All experiments related to *Mtb* H37Rv were conducted in the BSL-3 facility of CSIR-IMTECH.

#### Primary cell isolation

Peritoneal and Bone marrow derived macrophages were isolated exactly as described previously from C57BL/6 mice [18]. Adherent monolayer cells were confirmed to be pure macrophages by staining with CD11b antibody followed by flow cytometry (FACS Verse, BD Biosciences).

#### Cell culture, transfection, treatment and infection

J774 (mouse macrophage cell line) and RAW 264.7 cells were obtained from ECACC and National Centre for Cell Sciences (NCCS), Pune respectively and cultured in DMEM high glucose (Sigma) with 10% fetal bovine serum at 37°C under 5 % CO_2._ THP-1 cells were obtained from the National Centre for Cell Sciences (NCCS), Pune, India. Cells were cultured in RPMI-1640 supplemented with 10% fetal bovine serum. Peritoneal macrophages were isolated exactly as described previously from C57BL/6 mice. THP-1 cells were activated for 24 h with 25 ng/ml of phorbol 12-myristate 13 acetate (PMA), followed by 24 h of rest before further stimulation or infection. Transient transfections with siRNA duplexes were performed using Lipofectamine 2000 according to manufacturer’s instructions. Cells were analyzed 48 h after transfection. The mission esiRNA for SIRT2 (EMU017211), GAPDH (EMU184491) and Negative control (SIC001) were obtained from Sigma Aldrich.

To inhibit SIRT2 activity, 10 µM AGK2 (ab142073) have been utilized for all the experiments. For the infection experiments with AGK2, PMA-activated THP-1 cells were pre-treated with 10 µM AGK2 for 2 h following infection with *Mtb* at an MOI of 10. Clumping of bacterial cells was avoided by multiple passage of the cell suspension through a 26-gauge needle before infection in each experiment. This ensured that single bacterial cell suspensions were utilized for infection. Subsequently, cells were allowed to phagocytose *Mtb* for 4 h, after which non-phagocytosed bacteria were removed by washing the cells twice with serum free media. The infected cells were then maintained in complete RPMI-1640 along with 10 µM of AGK2 throughout the course of experiment. A parallel set of uninfected cells were also maintained to serve as control.

### Silencing of SIRT2

For cellular Sirtuin2 knockdown, THP-1 and J774 cells were transfected with human Sirtuin 2 short hairpin RNA (shRNA) lentiviral particles (Sigma-Aldrich) as per manufacturer’s instructions. To serve as control, separate sets of cells were transfected with pLKO.1-puro non-target shRNA control lentiviral particles (Sigma Aldrich). Stably transfected cells were selected and cultured in medium supplemented with puromycin (selection pressure of 7 µg/ml was utilized from 2 mg/ml puromycin stock for both THP-1 and J774 cells). Knockdown was confirmed by western blot of whole cell lysates using monoclonal mouse anti-SIRT2 antibody (ab211033). SIRT2 knockdown was confirmed using real-time quantitative PCR.

### Conjugation of Transferrin (Tf) / Apotransferrin (Apo Tf) with fluorophores

Tf and ApoTf were conjugated to Alexa 647 (Molecular probes) by reacting it for 1 hour at room temperature in 0.1 M sodium carbonate buffer (pH 9.0). All conjugates were purified or desalted by passing through buffer-equilibrated desalting column at 4°C.

### Analysis of surface receptor expression

THP-1 cells were infected with mid log-phase *Mtb* H37Rv GFP as described above. The cell surface expression of GAPDH, CD71 (TfR1), Holo-transferrin (Holo Tf), Apotransferrin (ApoTf) were assessed essentially as described previously. Briefly, cells were harvested in PBS-EDTA, washed with FACS buffer (phosphate buffer and 5% fetal bovine serum) and then blocked with FACS block (FACS buffer supplemented with 5% normal human serum and normal goat serum) at 4 °C for 30 min. Following blocking, cells were incubated with either 1 μg rabbit anti-GAPDH (Sigma, G9545) or its rabbit IgG isotype control (Invitrogen, 10500 C) for 1 h at 4 °C. The cells were then washed with FACS buffer three times and the bound anti-GAPDH antibody was detected with anti-rabbit Alexa 647 (Invitrogen, A21072). For surface CD71 staining, after blocking of cells, anti CD71-APC (BD pharmingen, 551374) or APC mouse IgG2a, k Isotype control (BD pharmingen, 555576), was used with 1 h incubation at 4 °C. To analyse the surface holotransferrin (holoTf) and Apotransferrin (ApoTf) binding, harvested cells were first blocked with 2% BSA at 4 °C for 30 minutes followed by incubation with 10 µg Tf-AF647 and Apo Tf-AF647 per tube at 4°C for 1 hour. Finally, cells were fixed in 4% paraformaldehyde or resuspended in PBS and analyzed using FACS Verse, FACS accuri or FACS Aria III Flow Cytometers (BD).

### Colony forming unit assay

PMA-activated THP-1 Empty vector and SIRT2 KD cells as well as Bone marrow derived macrophages (BMDMs) treated with or without AGK2 (10µM) were infected with *Mtb* H37Rv (at multiplicity of infection 1:10). Macrophages were lysed at 0, 24h and 48h with 0.05% SDS, serially diluted, and plated on Middlebrook 7H11 (Becton Dickinson, Difco, 212203) plates supplemented with 10% OADC (NaCl [Thermo fischer scientific, 15915], Dextrose [Thermo fischer scientific, Q24418], BSA fraction V [HIMEDIA,GRM105], Catalase [Sigma Aldrich,C40] and Oleic acid [Sigma Aldrich, 4954]). The plates were incubated at 37°C, and the *Mtb* colonies were counted after 3 weeks.

### Biotinylation of ApoTf, plasma membrane fraction isolation and Co-IP

Biotinylated ApoTf was prepared using Sulfo-NHS-LC-Biotin as per manufacturer’s instructions (Pierce). Biotin conjugated Apo Tf was purified and extensively washed with chilled PBS. 2×10^7^ Empty vector (EV) and SIRT2 knock down THP-1 cells were incubated with 500 µg of biotinylated Apo Tf in 1ml FACS buffer for 1 hour on ice. Controls were set in parallel wherein incubation of cells with biotinylated ApoTf was omitted. Cells were washed and processed for preparation of the membrane protein fractions. Membrane protein fractions were prepared as described previously [7]. Briefly, cells were homogenized in the homogenization buffer and the nuclear fraction was removed by centrifugation for 10 min at 700g at 4°C while the supernatant (S1) was collected and kept aside. The nuclear pellet was re-homogenized and spun as above. The supernatant (S2) was collected, and the final pellet was discarded. S1 and S2 were combined and centrifuged at 100,000g for 1 hour at 4°C. The resultant pellet was solubilized in phosphate buffer containing 0.2% Triton X-100 protease inhibitor mixture and agitated overnight at 4 °C to obtain the membrane protein fraction. The fractions were subjected to co-immuno precipitation using Streptavidin Magna beads (Pierce, Invitrogen) as per the manufacturer’s instructions. Beads were boiled in sample buffer and eluted proteins were analysed by western blot using monoclonal anti-GAPDH antibody.

### Western Blot

After infection or treatment with AGK2 as described in the respective legends, cells were washed with PBS and their lysates were prepared in RIPA cell lysis buffer (Gbiosciences, 786-70) along with protease inhibitor cocktail (Merck Millipore). The protein concentration was estimated using the BCA kit (Thermo scientific, 23227). Cell membrane fractions, whole cell lysates or Co-IP samples were resolved using 12% SDS-PAGE, transblotted onto PVDF membrane (Merck Millipore, IPVH00010) and blocked using 2% skim milk or 2% BSA. Blots were then probed with primary antibodies at 4°C overnight. Bound antibodies were then detected using HRP-linked secondary antibodies and developed using Luminata^TM^ Forte Western HRP Substrate (Merck Millipore WBLUF0500).

### Antibodies and reagents

Fth1 (3998) and Fpn1 (PA5-22993) antibodies were purchased from Cell Signaling Technology (CST) and Thermo scientific, respectively. The antibodies for Acetylated tubulin (T7451), β-actin (A2228), as well as anti-mouse (A4416) or anti-rabbit peroxidase antibody (A6154) were from Sigma. Antibody for SIRT2 (ab211033), SIRT2 inhibitor, AGK2 (ab142073) were obtained from Abcam. Antibody for GAPDH (CB1001) was from Calbiochem. Alexa fluor 568 and Alexa fluor 647 labelled secondary antibodies for confocal microscopy and flow cytometry were from Molecular probes, Invitrogen.

#### Analysis of labile iron pool by Calcein quenching assay

PMA-activated Empty vector (EV) and SIRT2 KD THP-1 or AGK2 (10µM) treated cells were washed three times with serum-free medium (SFM) incubated with 500 nM Calcein-AM (Invitrogen, C1430)) at 37°C for 15 minutes. Excess Calcein-AM was removed by extensive washing with SFM. Fluorescence of intracellular Calcein-AM was measured by flow cytometry (FACS Verse, BD bioscience) and the data were analyzed with FloweJo software. As Calcein fluorescence is quenched by iron, a decrease in cellular fluorescence is indicative of an increase in the intracellular labile iron pool.

### Real-Time PCR Analysis

Total cellular RNA from the uninfected and infected cells were isolated using TRIzol Reagent (Life Technologies) as per the manufacturer’s instruction. Synthesis of cDNA was carried out using RevertAid First Strand cDNA Synthesis Kit (Thermo Scientific #K1621) as per the manufacturer’s instruction followed by RT-PCR (Maxima SYBR Green master mix, #K0221) as per standard conditions using the gene specific primers. Briefly, the reaction mixture containing cDNA template, primers and maxima SYBR Green qPCR Master Mix was run in ABI Fast Real-time PCR System (Applied Biosystems, Foster City, CA, USA). Fold changes in mRNA levels were determined after normalization to internal control β-actin levels and analyzed using the 2C (T) method. The forward and reverse primers of utilized were designed with Primer-BLAST software and are listed as: Human SIRT2 Forward: 5՛-GAGGCCAGGACAACAGAGAG-3՛, Human SIRT2 Reverse: 5-՛GAGATTGGGGGATGTTCTGA-3՛, Human β-actin Forward: 5՛-TTCTACAATGAGCTGCGTGTG-3՛, Human β-actin Reverse: 5՛-GGGGTGTTGAAGGTCTCAAA-3՛

### 2D-PAGE analysis

Membrane proteins (500 µg from SIRT2 KD and high iron-treated cells) were processed by BioRad protein clean up kit® (163-2140) as per manufacturer’s instructions and subsequently the obtained pellet was dissolved in 125 µl of 2D-rehydration/sample buffer (BioRad, 163-2106). IPG 7 cm, pH 3-10 linear gradient strips (BioRad) were loaded with rehydrated sample solution. Isoelectric focussing was performed at 0-250 V for 2 h (linear), 250 V for 1 h (rapid), 250-3000 V for 4 h (linear), 3000 V-15,000V for 5 h. The current was limited to 50 µA per strip, and the temperature was maintained at 20°C for all isoelectric focussing steps. For the second dimension SDS-PAGE, the IPG strips were incubated in equilibration buffer I (BioRad, 163-2107) for 10 min followed by equilibration buffer II (BioRad, 163-2108) and then transferred onto 4-15% gradient acrylamide gels (BioRad). The gels were run at 25 mA until the bromophenol blue front had reached the bottom of the gel. Finally, gels were transferred to PVDF membrane and immunoblotted with anti-GAPDH antibody as described previously [19]. To ensure reproducibility, samples were analyzed in duplicate.

### Total iron estimation

Empty vector and SIRT2 KD THP1 cells (5 × 10^6^ ) either infected or uninfected were homogenized with “Iron Assay buffer” provided in the Fe estimation kit (Sigma, MAK025) and centrifuged at 16,000 × g for 10 minutes at 4 °C to remove the insoluble material. To measure total Fe, 50 µl of sample was added to 96-well plate and volume was adjusted to 100 µl with assay buffer. 5 µl of iron reducer was added to each of the sample wells. The plate was then mixed well using a horizontal shaker and incubated the reaction for 30 minutes at 25 °C while being protected from light. 100 μL of Iron Probe was added to each well containing standard and test samples. Contents in plate were mixed well using a horizontal shaker and the reaction was incubated for 60 minutes at 25 °C. The plate was protected from light during the incubation and the absorbance was measured at 593 nm (A_593_) as described in earlier reports [20].

### Co-localisation of cell surface GAPDH and Apo Tf

5×10^5^ THP-1 cells were cultured in confocal imaging dishes and activated for 24 hours with 25 ng/ml of phorbol 12-myristate 13-acetate (PMA). Subsequently the cells were allowed to rest for another 24 hours in fresh complete RPMI media. Cells were blocked with 2% BSA in SFM at 4°C for 30 minutes. Subsequently, cells were stained with 1 µg/ µl of mouse anti-GAPDH antibody (Calbiochem) in 100 µl FACS buffer for 1 hour followed by extensive washing. The anti-GAPDH antibody was subsequently detected using Alexa Fluor 568 (at dilution 1:1200, Invitrogen). Simultaneously, cells were stained with Apo Tf-FITC conjugate. After three washes, cells were fixed in 4% paraformaldehyde and imaged on Nikon A1R confocal microscope using a 60× oil immersion objective and 1 Airy unit aperture.

### Radioactivity

THP-1 macrophages were plated into 12 well plates (2× 10^5^ cells/well), PMA stimulated for adherence, rested for 24h to differentiate Subsequently, 100 nM Fe^55^ (ARC USA, #ARX0109) was added to the medium and after 24h of incubation with radioactive iron, cells were treated with AGK2 for 24h. Untreated cells were set up in parallel. Cells were washed extensively, and the radioactivity incorporated into cells was measured by liquid scintillation in the β-counter. Blank values (typically <100 cpm) were subtracted from all readings.

## Acknowledgements

Mr. Anil Theophilus & Mr. Randeep Sharma are acknowledged for technical assistance. We thank Dr. Neeraj Khatri and members of the IMTech Centre for Animal Resources and Experimentation for their assistance in obtaining clearance of projects form the Animal Ethics Committee. This is IMTECH communication no. 057/2023.

## Author contributions

MR and CIR conceptualized and finalized the manuscript. ST designed the study, investigated, collected data, performed analysis, and wrote the initial draft of paper. RM helped in BSL3-related flow cytometry experiments. AD assisted in cell culture and microscopy-related experiments. GKC and RD helped in the animal handling and isolation of primary cells.

## Funding

ST, AD and GKC received fellowships from Department of Biotechnology, India. RM received fellowship from University Grants Commission, India and RD was supported by fellowship from Indian Council of Medical Research. Financial support of; DST, DBT, CSIR, ICMR and UGC Government of India, by way of research grants and fellowships is acknowledged.

## Declarations

### Conflict of interest

The authors have no conflicts of interest to declare that are relevant to the content of this article.

## Notes

### Competing Interest Statement

The authors have declared no competing interest.

