## Supplemental Information for "*Mycobacterium tuberculosis* exploits SIRT2 for iron acquisition to facilitate its intracellular survival"

**Supplementary Figure S1A**

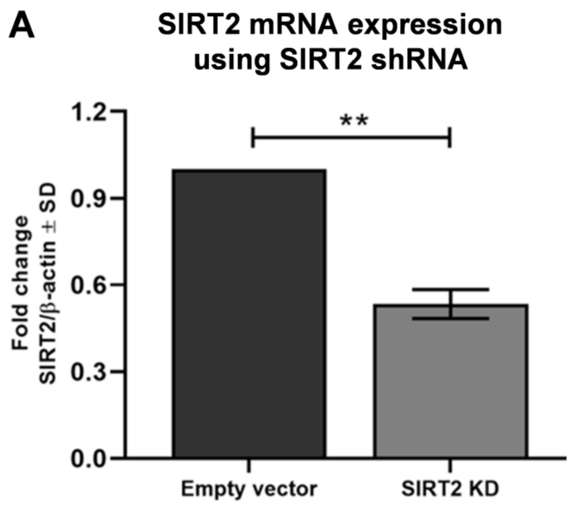


**Fig. S1A** mRNA levels of SIRT2 in Empty vector (EV) and SIRT2 KD THP1 macrophages assessed by RT-PCR (n=3, **p<0.01).

**Supplementary Figure S2**

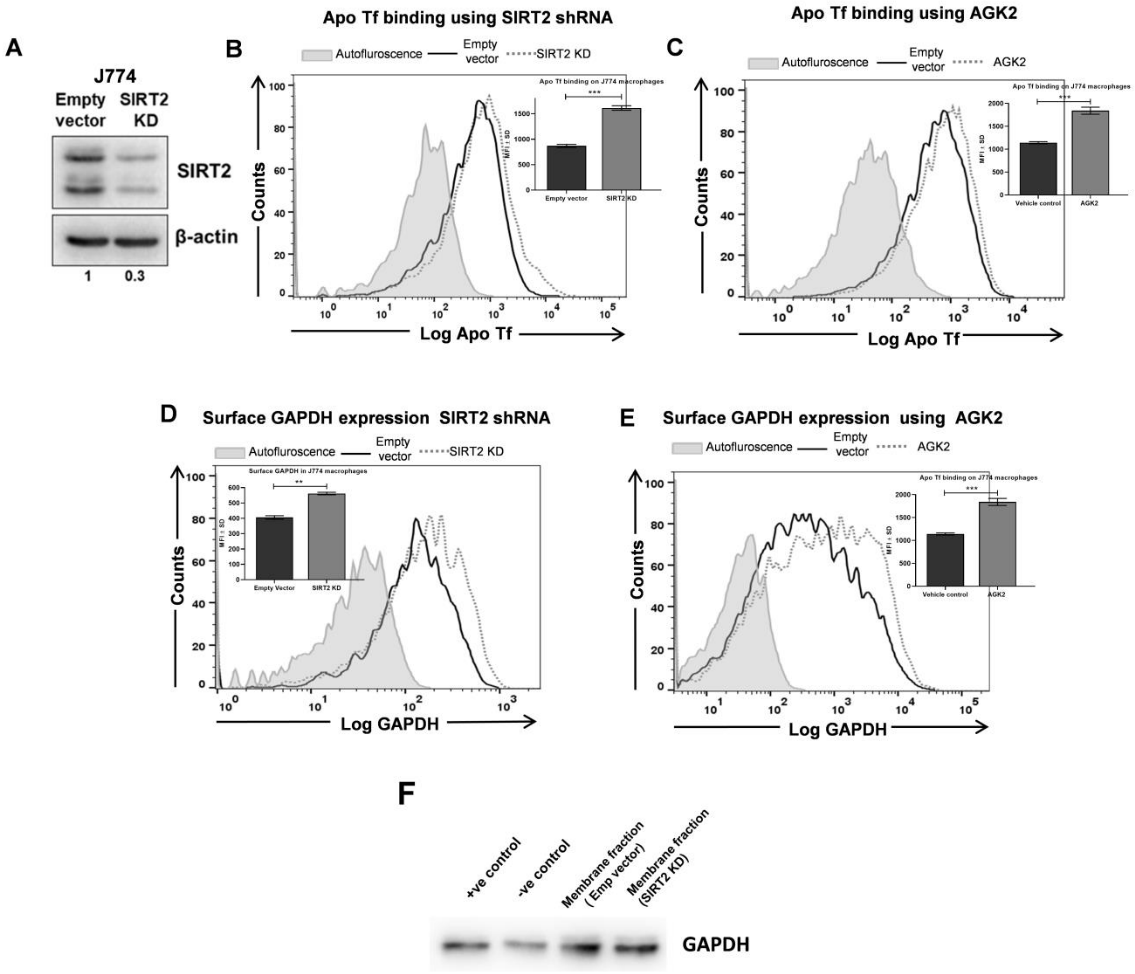


**Fig S2 SIRT2 up regulates the expression and binding of GAPDH and Apo Tf in J774 macrophages (A)** Representative immunoblot of SIRT2 in EV and SIRT2 KD J774 macrophages for confirmation of SIRT2 KD **(B-C)** Apo Tf binding in EV/ Control and SIRT2 KD/ AGK2 macrophages after incubation of labelled Apo Tf at 4$^{o}$C for 1h. Representative flow cytometry overlay has been shown, data in inset are presented as representative plot of background corrected mean fluorescence intensity (MFI ± SD) from 2 independent experiments performed in triplicates (n=3, ***p<0.001) **(D-E)** EV/Control and SIRT2 KD/ AGK2-treated macrophages were incubated with anti-GAPDH antibody for quantification by flow cytometry**.** Representative flow cytometry overlay has been shown, data in inset are presented as representative plot of background corrected mean fluorescence intensity (MFI ± SD) from 2 independent experiments performed in triplicates (n=3, ***p<0.001) **(F)** EV and SIRT2 KD THP1 macrophages were incubated with biotinylated apotransferrin at 4˚C followed by plasma membrane fraction preparation and subjected to co-immunoprecipitation (Co-IP) using streptavidin–Magnabeads. Using monoclonal anti-GAPDH antibody, GAPDH was detected in both the fractions suggesting an interaction between GAPDH and Apo Tf. A negative (-ve) control was run in parallel, wherein the incubation of cells with biotinylated apotransferrin was omitted. Standard rabbit muscle GAPDH was used as a Positive (+ve) control.

**Supplementary Figure S3**

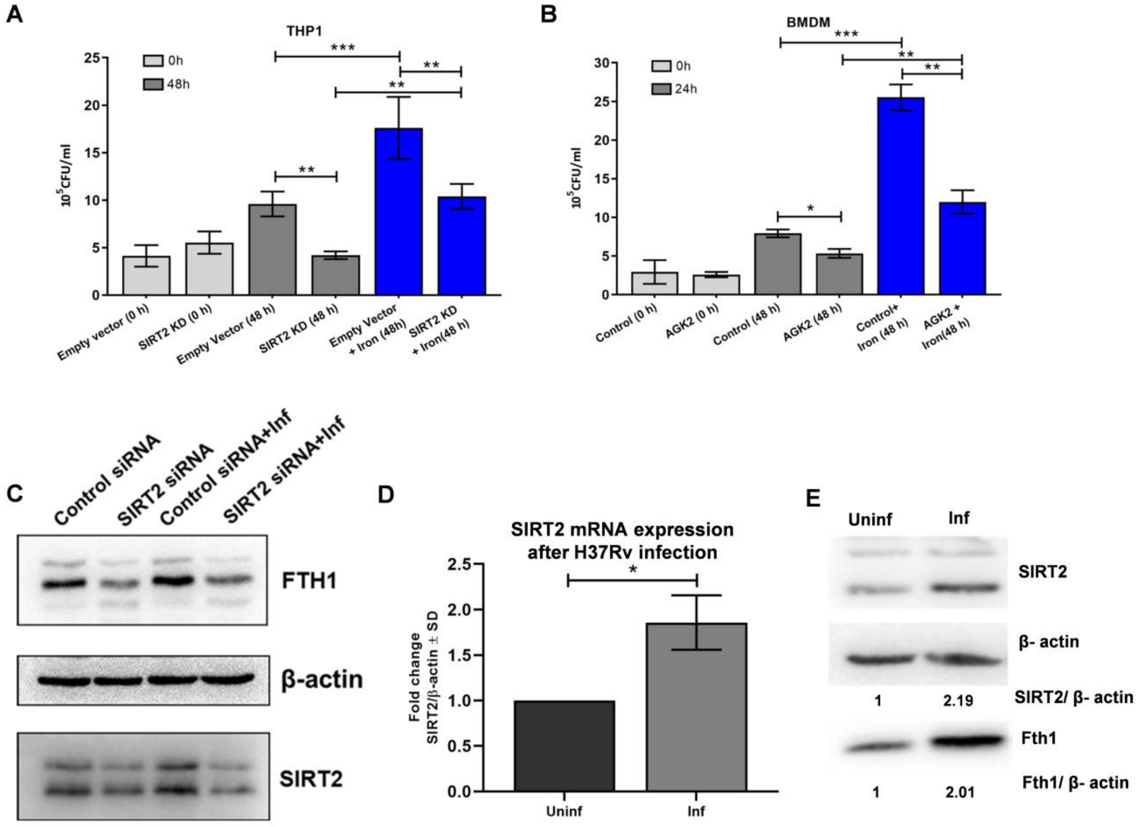


**Fig. S3 (A)** Empty vector and SIRT2 KD THP-1 macrophages were infected with 1:10 MOI of *Mtb* H37Rv and 100 µM $FeCl_3$ was added wherever appropriate. Cells were processed for CFU enumeration at 0 h and 48 h hours post infection. Results are indicated as $10^5$ CFU/mL (***p<0.0001 and **p<0.01, *p<0.05, n=3). Data represents average of two independent experiments performed in triplicates **(B)** Bone marrow derived macrophages were treated with or without AGK2 for 24 h followed by infection with 1:10 MOI of *Mtb* H37Rv and 100 µM $FeCl_3$ wherever appropriate. After certain time points, CFU load was measured. Results are indicated as $10^5$ CFU/mL (***p<0.0001 and **p<0.01, *p<0.05, n=3). Data represents average of two independent experiments performed in triplicates **(C)** Representative blot showing expression of Fth1 with or without H37Rv infection in control siRNA and SIRT2 siRNA transfected THP1 macrophages **(D)** RT-PCR analysis of SIRT2 in macrophages with or without infection of *Mtb* for 24 h at 1: 10 MOI **(E)** Western blot of SIRT2 in peritoneal macrophages with or without infection of *Mtb* for 24 h at 1: 10 MOI.
